# Inline Liquid Chromatography-Fast Photochemical Oxidation of Proteins Allows for Targeted Structural Analysis of Conformationally Heterogeneous Mixtures

**DOI:** 10.1101/825521

**Authors:** Surendar Tadi, Sandeep K. Misra, Joshua S. Sharp

## Abstract

Structural analysis of proteins in a conformationally heterogeneous mixture has long been a difficult problem in structural biology. In structural analysis by covalent labeling mass spectrometry, conformational heterogeneity results in data reflecting a weighted average of all conformers, complicating data analysis and potentially causing misinterpretation of results. Here, we describe a method coupling size exclusion chromatography (SEC) with hydroxyl radical protein footprinting using inline fast photochemical oxidation of proteins (FPOP). Using a controlled synthetic mixture of holomyoglobin and apomyoglobin, we demonstrate that we can achieve accurate footprints of each conformer using LC-FPOP when compared to off-line FPOP of each pure conformer. We then applied LC-FPOP to analyze the adalimumab heat-shock aggregation process. We found that the LC-FPOP footprint of unaggregated adalimumab was consistent with a previously published footprint of the native IgG. The LC-FPOP footprint of the aggregation product indicated that heat shock aggregation primarily protected the hinge region, suggesting this region is involved with the heat shock aggregation process of this molecule. LC-FPOP offers a new method to probe dynamic conformationally heterogeneous mixtures such as biopharmaceutical aggregates, and obtain accurate information on the topography of each conformer.

## Introduction

Hydroxyl radical protein footprinting (HRPF) is a method for probing the topography of proteins in solution. In HRPF, a protein of interest is exposed to hydroxyl radicals freely diffusing in solution. One method of generating radicals for HRPF, fast photochemical oxidation of proteins (FPOP) generates hydroxyl radicals in a flow system by the photolysis of hydrogen peroxide using a pulsed UV laser. The flow rate of the sample is matched to the pulse rate of the laser to ensure that each volume of sample is illuminated by a single ~20 ns laser pulse, preventing artifactual labeling^1–2^. These radicals are short-lived, highly reactive, and rapidly oxidize amino acid side chains present on the surface of the folded protein. The apparent rate of oxidation is primarily a function of two factors: the inherent chemical reactivity of the amino acid side chain^3–4^ and the exposure of the side chain to the hydroxyl radical, which correlates directly with solvent accessible surface area^5–7^. Information about the topography of the folded protein is frozen in this chemical “snapshot” of the protein surface, after which the protein can be processed (e.g. deglycosylated, extracted from lipid membranes, proteolytically digested, etc.) and analyzed by liquid chromatography coupled to mass spectrometry (LC-MS). By comparing the signal intensity from an unoxidized peptide versus all oxidized forms of that peptide, the apparent rate of oxidation for a particular peptide can be determined. HRPF is typically used to compare the same or very closely related amino acid sequences in two or more conformations. In such experiments, the inherent reactivity of a given amino acid side chain does not differ between samples, so changes in the apparent reactivity of an amino acid side chain due to hydroxyl radicals can be directly correlated with changes in the solvent-accessible surface area^6, 8–10^.

Systems where HRPF has found success in probing protein structure and protein interactions include membrane proteins^11–17^, protein oligomerization and aggregation processes^18–20^, and binding interfaces between proteins and unpurified complex mixtures of ligands^21^. What many of these difficult biological systems have in common is the presence of more than a single structure in solution to probe, which we will refer to here as conformational heterogeneity. Conformationally heterogeneous systems have the distinction of being both difficult to analyze with modern techniques as well as areas of intense interest due to an emerging understanding of their importance in biology and biochemistry. Standard high-resolution structural technologies have difficulties with conformational heterogeneity, as the presence of multiple major conformations can prevent protein crystallization, dampen signals in both crystallography and NMR, and make data very difficult to interpret. Conformational heterogeneity can arise from a single protein that samples multiple conformers in solution, post-translational modifications (PTMs) resulting in conformational changes, prosthetic groups, chemical modifications, co-existing sequence variants, or oligomerization and aggregation processes where the thermodynamics of protein aggregation and disaggregation results in a distribution of multimeric structures with topographies differing based on the number and arrangement of subunits. In each case of conformational heterogeneity, multiple topographies of the same protein sequence exist simultaneously in solution, with each topography playing an important role in the observed function (or dysfunction, as the case may be) of the protein. While in some cases these conformationally heterogeneous samples can be separated chromatographically for individual analysis of each conformer, if the heterogeneity is dynamic (e.g. aggregation processes) the separated sample may not remain conformationally homogeneous long enough for structural interrogation.

Here, we describe the development of inline LC-FPOP combining size exclusion chromatography (SEC) to separate conformationally heterogeneous mixtures and FPOP to individually footprint each conformer as it elutes from the column **(Fig. 1)**. As FPOP makes a covalent “chemical snapshot” of the conformation at the time of labeling immediately postcolumn, any mixture that is sufficiently conformationally stable to resolve by SEC LC can be analyzed by this method, even if the conformation shifts after labeling. A mixture of holomyoglobin and apomyoglobin was used to validate the accuracy of LC-FPOP results. LC-FPOP was then used to describe the heat shock aggregation products of an adalimumab biosimilar, an anti-TNFα IgG monoclonal antibody. Using LC-FPOP, we are able to identify the regions of adalimumab with altered topography gaining insight into the mechanism of aggregation in this biosimilar.

**Fig. 1.**
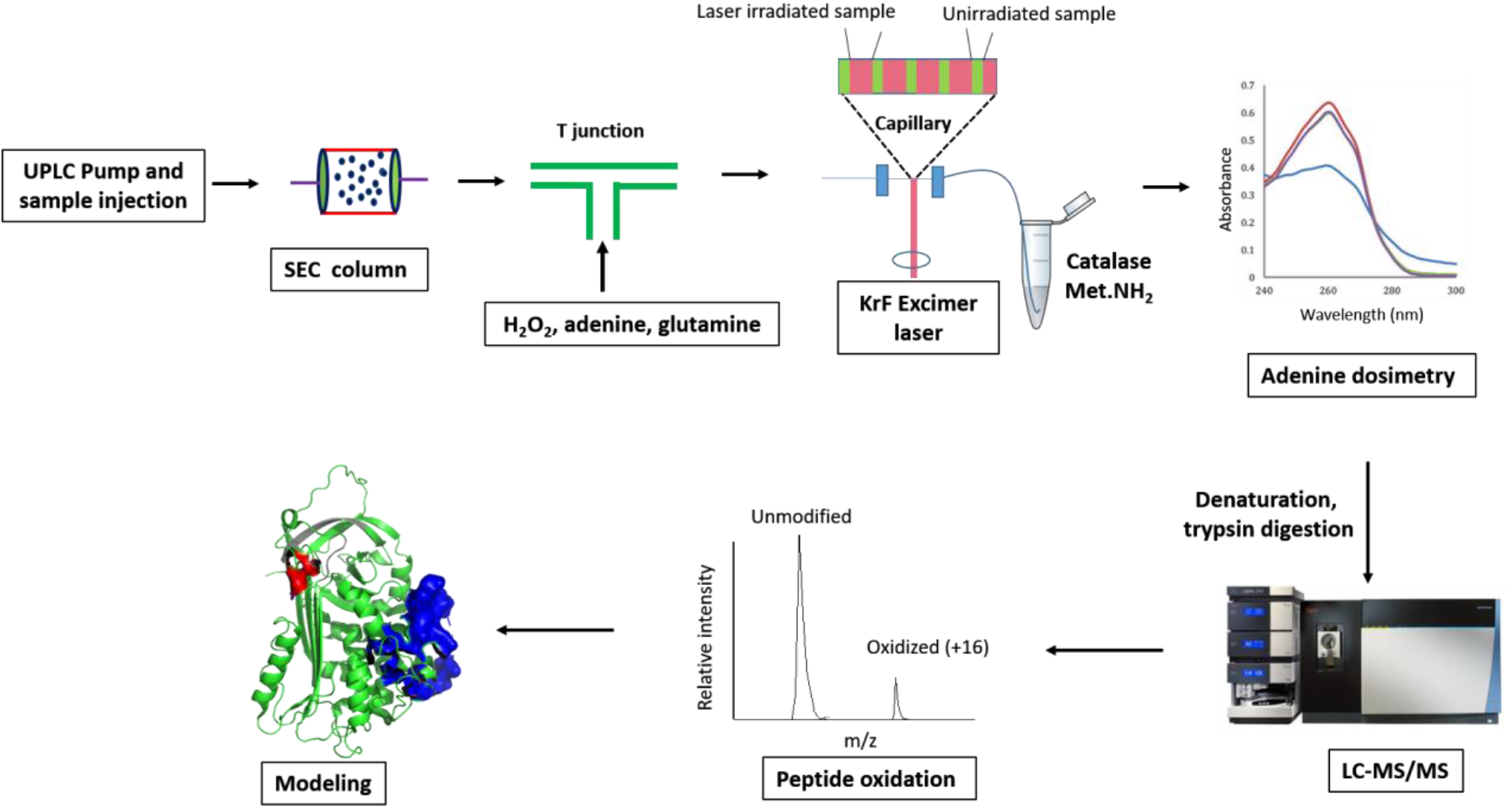
Schematic of inline FPOP. Protein mixture of heterogeneous conformations is separated on size exclusion chromatography. FPOP reagent is injected with T-junction, just before laser irradiation. FPOP reagent contains phosphate buffer, adenine and glutamine. After the laser irradiation, the samples are collected in a quench solution containing catalase and methionine amide. Adenine absorbance is measured to make sure that all replicates have similar amount of free radicals. After the protein digestion, the samples are run onto mass spectrometer, and footprint of the individual protein conformers are calculated and plotted onto protein structure.

## Results

### SEC separation and Inline FPOP of holomyoglobin and apomyoglobin

A mixture of holomyoglobin and apomyoglobin was used to test the ability to generate accurate HRPF data from a single polypeptide chain in two distinct conformations that can be readily analyzed in each separate state for validation. We developed a method for the isocratic aqueous separation of apomyoglobin and holomyoglobin using SEC chromatography. The injection of pure holomyoglobin gives one major peak that elutes around 42 minutes, while the injection of pure apomyoglobin gives one major peak around 65 minutes **(Extended Data Fig. 1a,b)**. SEC separation of a synthetic 1:1 mixture of holomyoglobin and apomyoglobin was easily achieved under these conditions **(Extended Data Fig. 1c)**. The baseline resolution of holomyoglobin and apomyoglobin indicates simple and efficient chromatographic separation between the two conformers of the same protein.

Sample corresponding to holomyoglobin and apomyoglobin were analyzed via inline SEC LC-FPOP as outlined in **Fig. 1**, and labeled sample was collected as guided by the UV trace from 35 to 45 min and 65 to 75 min respectively. Samples were immediately quenched upon collection after FPOP to eliminate hydrogen peroxide and any secondary oxidants ^1, 22^. Adenine dosimetry was used to ensure that samples were exposed to equivalent radical doses^23–24^. The results from inline LC-FPOP of a 1:1 mixture of holomyoglobin and apomyoglobin were compared with traditional off-line FPOP of each pure component to determine if inline LC-FPOP generated comparable results from a mixture of products. Results from LC-FPOP and traditional off-line FPOP were remarkably consistent with one another. Almost identical sequence coverage was obtained from samples, regardless if FPOP of the mixture was performed inline or off-line. Similarly, almost identical footprint pattern was obtained by inline LC-FPOP and off-line FPOP of the pure conformer mixture **(Fig. 2)**. Extracted ion chromatogram of the +16 Da oxidation products of the peptide 103-118 shows clear conformational differences between apomyoglobin and holomyoglobin. The LC trace shows a highly similar profile and relative quantity of oxidation products between the inline LC-FPOP of the mixture, and off-line FPOP of the pure conformer. Both inline and traditional methods generate very different oxidation profiles between apomyoglobin and holomyoglobin; however, the peptide oxidation products for off-line FPOP and inline FPOP are indistinguishable indicating LC-FPOP of a conformer recapitulates the data generated from traditional FPOP of a pure conformer **(Extended Data Fig. 2)**. These data indicate that LC-FPOP of a mixture allows the researcher to obtain equivalent topographical information as traditional off-line FPOP of the pure conformers.

**Fig. 2.**
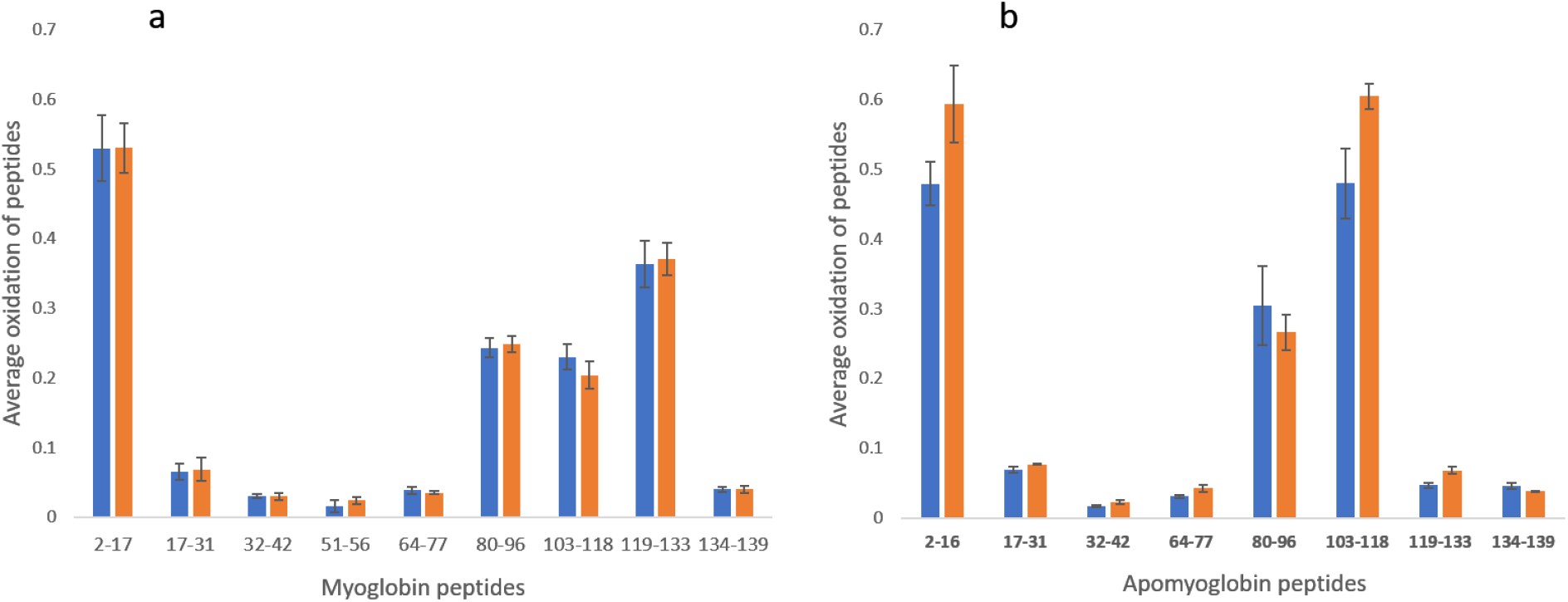
Comparison of inline LC-FPOP (blue) and off-line FPOP (orange). The mixture of holomyoglobin and apomyoglobin was separated using SEC. Comparison of peptide oxidation between FPOP performed inline and off-line of holomyoglobin **(a)** and apomyoglobin **(b)**. Error bars represent one standard deviation from a triplicate data set. The footprint pattern was almost identical between the inline and off-line FPOP.

### SEC separation and FPOP of native and heat-shock aggregated adalimumab biosimilar

Inline FPOP was used to measure the aggregate of the pharmacologically important biosimilar monoclonal antibody adalimumab. Injection of monomeric adalimumab gives one major peak that elutes around 30 minutes. We heated the adalimumab at 80 °C for 15 min to initiate the aggregation process. The injection of the heat shocked adalimumab results in a single aggregate peak around 25 min. SEC separation of a 1:1 mixture of monomeric adalimumab and aggregated adalimumab was easily achieved under these SEC conditions **(Fig. 3)**.

**Fig. 3.**
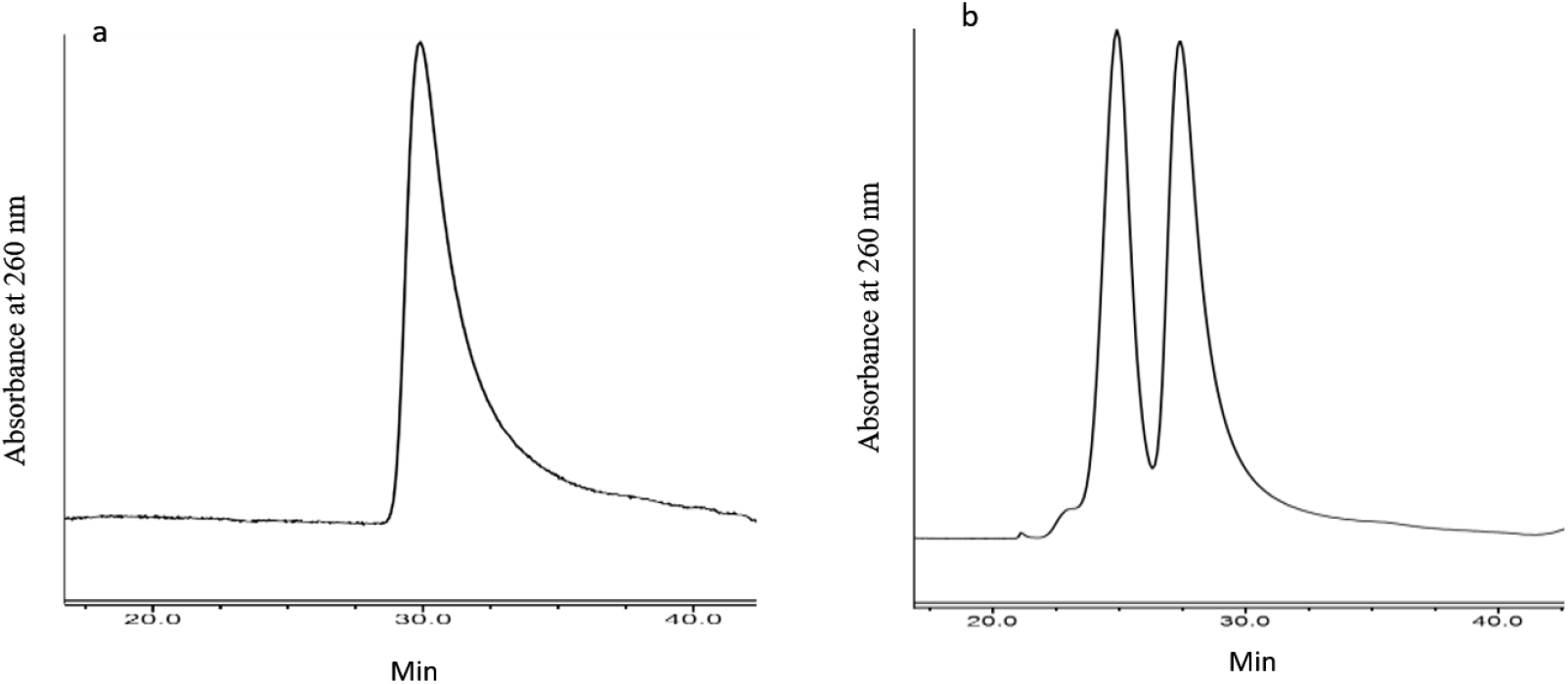
Size exclusion chromatography of adalimumab. **a)** The native adalimumab gives a single peak around 30 min. The adalimumab was heated at 80 °C for 15 min to make aggregates. **b)** Separation of monomeric and aggregated adalimumab by SEC.

We then performed inline FPOP of the native and aggregated adalimumab (DLS characterization shown in **Extended Data Fig. 3**). A mixture of 5 mg/mL native and aggregated adalimumab was run through SEC column. The FPOP was performed as proteins eluted from the column as described previously. The sample after the laser irradiation was collected and quenched separately corresponding to the native and aggregated adalimumab. Monomeric adalimumab comprised the fraction collected from 26 to 30 min and aggregated adalimumab correspond to the fraction between 22 to 26 min. Comparison of monomeric adalimumab with previously published FPOP results^24^ showed that the LC-FPOP results of monomeric adalimumab obtained from the mixture and previously published traditional FPOP results are highly comparable (**Extended Data Fig. 4)**.

Comparison of the footprint for the peptides from native and aggregated adalimumab was carried out to determine the regions of structure impacted by aggregation **(Fig. 4a)**. As expected, there are significant differences in the peptide oxidation values between the native and aggregated adalimumab for a large number of peptides from both the heavy as well as light chains. When the results of this FPOP experiment are plotted on the three-dimensional structure of adalimumab **(Fig. 4b)**, the spatial organization of the different areas of exposure and protection allows for interpretation. The results suggest that the hinge region of adalimumab experiences the most widespread protection upon heat shock aggregation, with little change in the end of the Fc region or for much of the paratope. Indeed, some paratope-adjacent portions of the protein experience modest exposure upon aggregation. Almost all of the light chain experience a significant change in oxidation, with all but the N-terminal peptide showing significant protection by aggregation. By contrast, the end of the Fc region is largely unchanged, indicating that the hinge region and adjacent areas mediate the heat shock-induced aggregation of adalimumab.

**Fig. 4.**
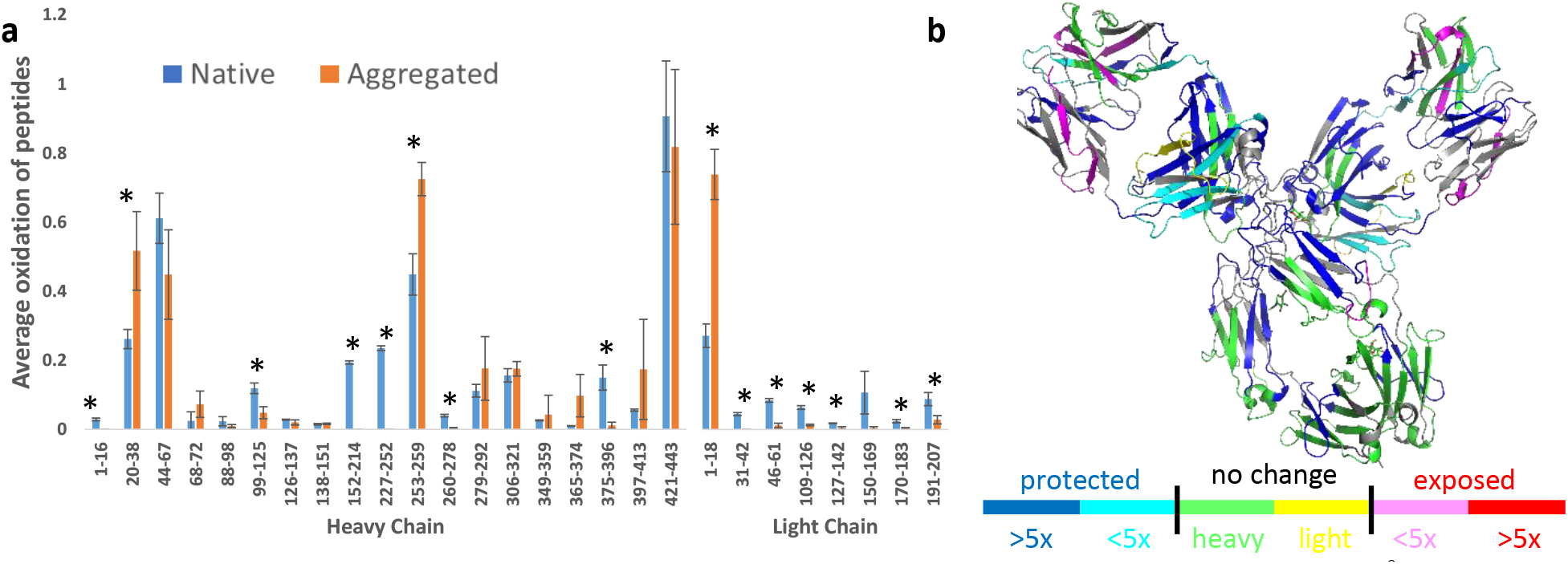
FPOP of the native and aggregated adalimumab. Adalimumab heated at 80 °C for 15 min results in its aggregation. A mixture of monomer and aggregated adalimumab was separated on SEC followed directly by in-line FPOP of the monomer and aggregate peaks. **(a)** The comparison of the footprint of (left) heavy chain and (right) light chain peptides monomer (blue) and aggregated (orange) adalimumab. Error bars represent one standard deviation from triplicate measurements. Asterisks label peptides that significantly change in oxidation between the monomer and aggregate (*p* ≤ 0.05). **(b)** LC-FPOP data are mapped onto a model of adalimumab. The model is colored to represent the change in oxidation in aggregate compared to monomer. Blue: >5x decrease; cyan: <5x decrease; magenta: <5x increase; green: no change (heavy chain); yellow: no change (light chain); grey: peptide not observed.

## Discussion

Hydroxyl radical protein footprinting as a technology has proven to have tremendous flexibility, offering structural insights into highly challenging systems including oxidation-induced conformational change^9, 25^, observing structural waters in integral membrane proteins^14, 16, 26–27^, differentiating infectious from non-infectious prion aggregates^28^, mechanisms of allosteric inhibition^29^ and protein-carbohydrate interactions^6, 21, 30–31^, to name a few. However, in all cases, the resulting HRPF footprint represents the average of all conformers present in solution. The ability to individually interrogate each conformer previously required the ability to purify and prepare each conformer individually, which is not always possible or practical.Here, we demonstrate the use of inline LC-FPOP to probe individual conformer topography of mixtures of two conformers, with results that are comparable to traditional FPOP analysis of either pure conformer. We targeted conformers that are non-dynamic (apomyoglobin vs. holomyoglobin) to ensure proper control of conformer composition in our mixtures, then used the method to demonstrate analysis of conformers that can shift dynamically, but at a slow time scale at room temperature (monomeric vs. aggregated adalimumab). The ability to structurally differentiate non-dynamic conformers from a mixture is very convenient, especially for the structural characterization of complex analytes like disulfide bond shuffling products^32^. However, the ability of LC-FPOP to also work for dynamic conformer mixtures enables detailed analyses of new systems, so long as the dynamics are slow enough to achieve separation on a chromatographic time scale. These mixtures include a surprisingly broad array of biomedically important systems that are currently subjects of intense investigation, including not only monoclonal antibody aggregation^33^, but protein misfolding^34^, oligomerization of amyloids such as tau^35^, protein-polysaccharide complexes^36^ and slow-exchanging conformers in intrinsically disordered proteins^37^. Given the problems that exist in the structural analysis of these systems currently, LC-FPOP represents a significant new tool for enabling structural investigations.

We demonstrate LC-FPOP using SEC in order to achieve separation using an isocratic gradient with aqueous buffer. The LC-FPOP technology is not limited to SEC; any chromatography that uses an isocratic gradient could potentially be used for LC-FPOP so long as the buffer system is compatible with FPOP. An isocratic gradient is convenient in order to minimize changes in the hydroxyl radical scavenging capacity of the solvent. However, with the recent report of inline hydroxyl radical dosimetry^38^, binary solvent systems can be designed with matched radical scavenging capacities for aqueous separations (for example, a binary gradient of sodium formate/sodium chloride for strong anion exchange chromatography) and tested in real-time for suitability. The ability to couple inline radical dosimetry with a method for real-time control of hydroxyl radical generation would allow for real-time scavenging compensation^38^, providing even more flexibility in LC gradient design. As the use of adsorptive stationary phases and faster separation times would allow for probing of dynamic systems with more subtle changes in conformation and faster kinetics of conformational change, we are pursuing these advances in current studies.

## Materials and methods

### Materials

Adalimumab was obtained from GlycoScientific (Athens, Georgia). Catalase, glutamine, formic acid, hydrochloric acid, sodium phosphate, and 2-(N-morpholino)ethanesulfonic acid (MES) were purchased from Sigma-Aldrich (St. Louis, MO). Adenine and LC-MS grade acetonitrile and water were purchased from Fisher Scientific (Fair Lawn, NJ). Hydrogen peroxide (30%) was purchased from J. T. Baker (Phillipsburg, NJ). Fused silica capillary was purchased from Molex, LLC (Lisle, IL). Sequencing grade modified trypsin was obtained from Promega (Madison, WI).

### Inline FPOP

5 mg/mL adalimumab protein was heated at 80 °C for 15 min and cooled on ice immediately for 2 min to make aggregates. Following aggregation, 5 mg/mL of native and aggregated protein mixture was loaded on a size exclusion column (ACQUITY UPLC Protein BEH 150 mm, 125 Å, 1.7 μm, Waters, Milford, MA) using Dionex Ultimate 3000 (Dionex, Sunnyvale, CA). Separation of proteins was performed with an isocratic gradient of 100 mM sodium phosphate (pH 6.8) at the flow rate of 30 μL/min for 60 min. Column eluant flowed through a Dionex UV detector, where protein elution was detected by UV absorbance at 280 nm. A Peek microTee (Upchurch Scientific) mixer was installed immediately after the UV detector, with one inlet port leading to the SEC column, one inlet port to the FPOP reagent, and the outlet port to the fused silica capillary (365 μm O.D. 100 μm I.D.) for laser exposure. FPOP reagent was mixed 1:1 with the eluant using a Legato 101 syringe pump (KD Scientific, Holliston, MA) with a gastight syringe (Hamilton, Reno, NV), to a final concentration of 100 mM hydrogen peroxide, 16 mM glutamine, and 2 mM adenine. Adenine radical dosimetry is used to ensure comparable radical exposure of both components of the mixture.

After the mixing micoTee, the sample was passed through the focused beam path of a COMPex Pro 102 KrF excimer laser (Coherent Inc., Santa Clara, CA). The exclusion volume was set to 15%, the fluence was 13 mJ/mm^2^ and the frequency of the laser was set to 20 Hz. Samples were collected from 23 to 25 and 27 to 30 min immediately after illumination into vials containing 120 μL of a quench solution containing 0.5 μg/μL methionine amide and 0.2 μg/μL catalase to eliminate secondary oxidation. After quenching, the absorbance of the adenine dosimeter at 265 nm was measured using a Nanodrop UV/Vis spectrophotometer. Adenine dosimetry was used to ensure that samples were exposed to equivalent radical doses^23^. Samples were concentrated in a lyophilizer and were analyzed on a LC-MS/MS system.

### Off-line FPOP

All samples were prepared and analyzed in triplicate. A final concentration of 5 mg/mL each of holomyoglobin or apomyoglobin was mixed with a final concentration of 50 mM sodium phosphate pH 7.4, 1 mM adenine, 17 mM glutamine and 200 mM hydrogen peroxide were added just before the laser exposer loaded on gastight syringe (Hamilton, Reno, NV). The mixture was flowed through the focused beam path of the excimer laser pulsing at 20 Hz, with an exclusion volume 15% and a fluence of ~13 mJ/mm^2^. The exposed sample was collected into vials containing 0.3 μg/μL of catalase and 0.5 μg/mL of methionine amide^24, 38^. Samples were incubated at room temperature for 30 min and adenine absorbance was measured to ensure that all the samples received comparable amount of free radicals. Samples were processed for LC-MS analysis as described below.

### Sample Digestion

50 mM Tris, pH 8.0 and 1 mM CaCl2 was added to samples after inline FPOP. The samples were incubated at 90 °C for 15 min to denature the protein. After denaturation, samples were cooled to room temperature and a 1:20 trypsin/protein weight ratio was added to the samples for overnight digestion at 37 °C with sample mixing. Digestion was terminated by heating the samples to 95 °C for 10 min.

### LC-MS Analysis

The samples were loaded on to an Acclaim PepMap 100 C18 nanocolumn (0.75 mm × 150 mm, 2 μm, Thermo Fisher Scientific). Separation of peptides on the chromatographic system was performed using mobile phase A (0.1% formic acid in water) and mobile phase. B (0.1% formic acid in acetonitrile) at a flow rate of 300 nL/min. The peptides were eluted with a gradient consisting of 2 to 35% solvent B over 22 min, ramped to 95% solvent B over 5 min, held for 3 min, and then returned to 2% solvent B over 3 min and held for 9 min. Peptides were eluted directly into the nanospray source of an Orbitrap Fusion instrument controlled with Xcalibur version 2.0.7 (Thermo Fisher, San Jose, CA) using a conductive nanospray emitter (Thermo Scientific). All data were acquired in positive ion mode. The spray voltage was set to 2400 V, and ion transfer tube was set to 300 °C. In CID mode, full MS scans were acquired from m/z 350 to 2000 followed by eight subsequent MS/MS scans on the top eight most abundant peptide ions.

### FPOP data analysis

Data acquired by LC-MS/MS of oxidized and unoxidized peptides were initially identified by Byonic version v2.10.5 (Protein Metrics, San Carlos, CA) for protein sequence coverage analysis. Unoxidized and oxidized peptide peaks were quantified by integration of the selected ion chromatogram peaks of unoxidized and oxidized peptides respectively plus one or more oxygen atoms (mass error = 10 ppm), with all resolved oxidation isomers summed using Xcalibur. Oxidation events per peptide were calculated using eq 1

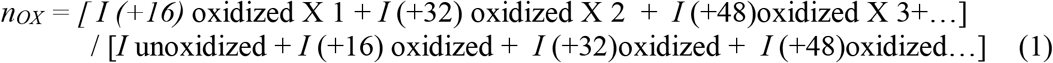

Oxidation events at peptide level were denoted as *n_OX_* and peak intensities of oxidized and unoxidized were denoted as *I*. All major identified oxidation products are the net addition of one or more oxygen atoms.

## Acknowledgments

This work was supported by the National Science Foundation (1608685) and the National Institute of General Medical Sciences (R01GM127267). J.S.S. and S.K.M. acknowledge support for LC-MS analysis from the Glycoscience Center of Research Excellence (NIH P20GM103460).

## Author contributions

S.T. performed LC-FPOP and traditional offline FPOP of all samples, performed data analysis of myoglobin samples, and wrote portions of the manuscript. S.K.M. analyzed LC-FPOP data for adalimumab, performed DLS analysis, and wrote portions of the manuscript. J.S.S. designed experiments, analyzed data for LC-FPOP, traditional FPOP and DLS, and wrote portions of the manuscript.

