## Extended Data for "Inline Liquid Chromatography-Fast Photochemical Oxidation of Proteins Allows for Targeted Structural Analysis of Conformationally Heterogeneous Mixtures"

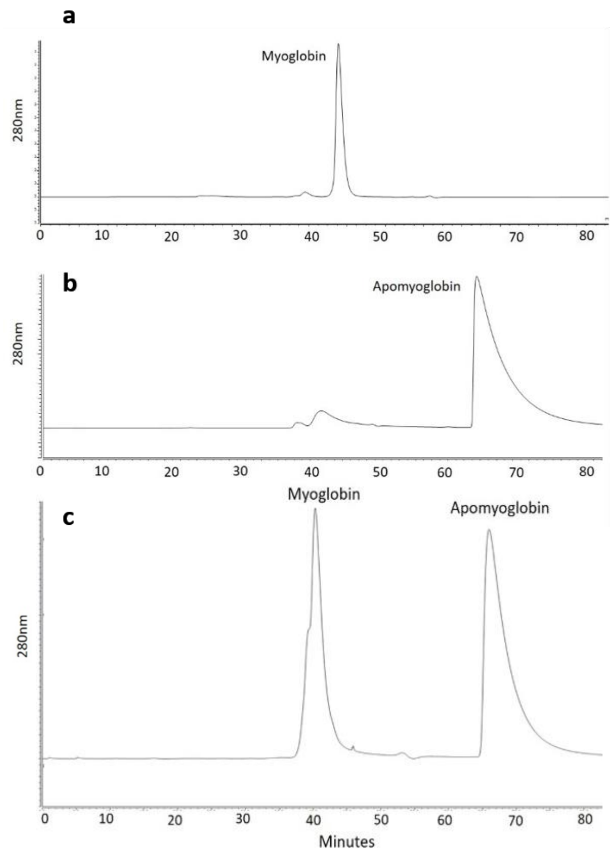


**Extended Data Fig. 1. SEC separation of holomyoglobin and apomyoglobin**. Injection of **(a)** only holomyoglobin results in a peak at 280 nm around 45 min and **(b)** only apomyoglobin results in peak detection around 68 min**.** The 1:1 mixture of holomyoglobin and apomyoglobin results in two distinct peaks, indicating the resolving power of the SEC used to separate the same protein with and without the heme.

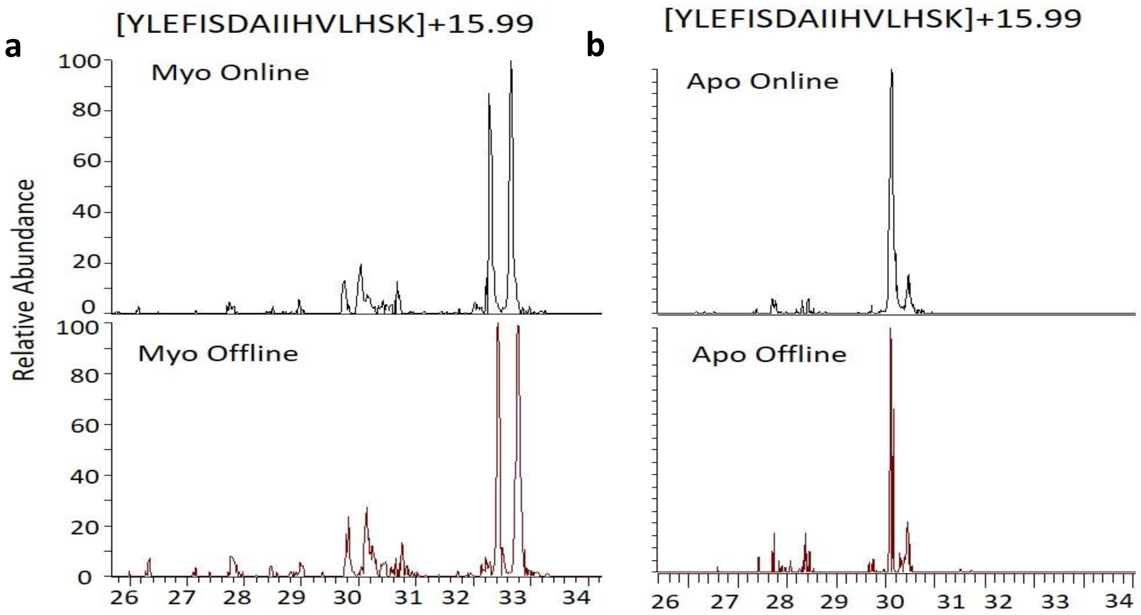


**Extended Data Fig. 2. Comparison of the extracted ion chromatogram**. Figure shows the +16 Da oxidation products of peptide 103-118 from **(a)** holomyoglobin and **(b)** apomyoglobin. The top panels in the figure is from online LC-FPOP of a 1:1 mixture, while the bottom panel is from traditional FPOP of the pure protein conformer.

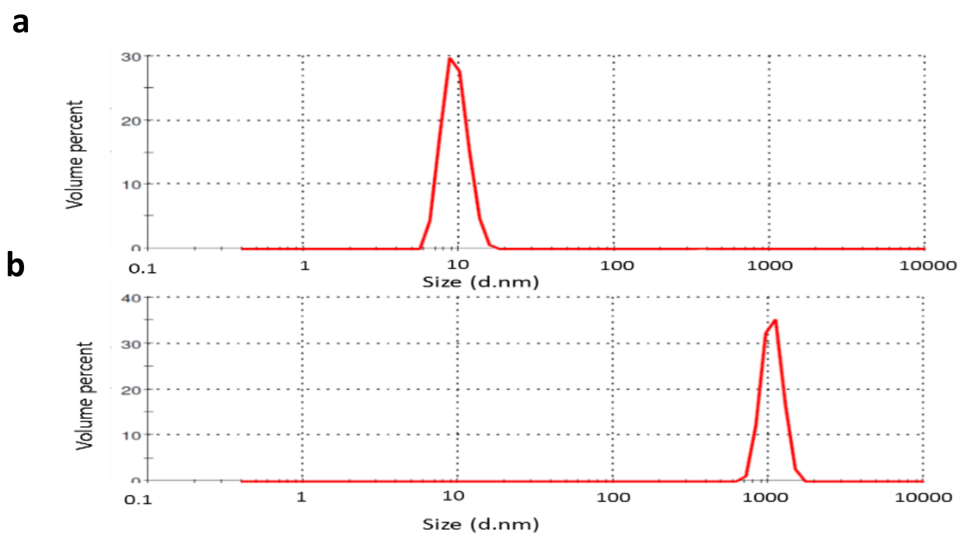


**Extended Data Fig. 3. Volume-mode dynamic light scattering of adalimumab**. (a) Native adalimumab has a Z-average particle size of about 9.5 ± 1.7 nm. When adalimumab was heated at 80 $^{0}$C for 15 min to make aggregates, the Z-average size of the particle is shifted to 954.9 ± 140.9 nm.

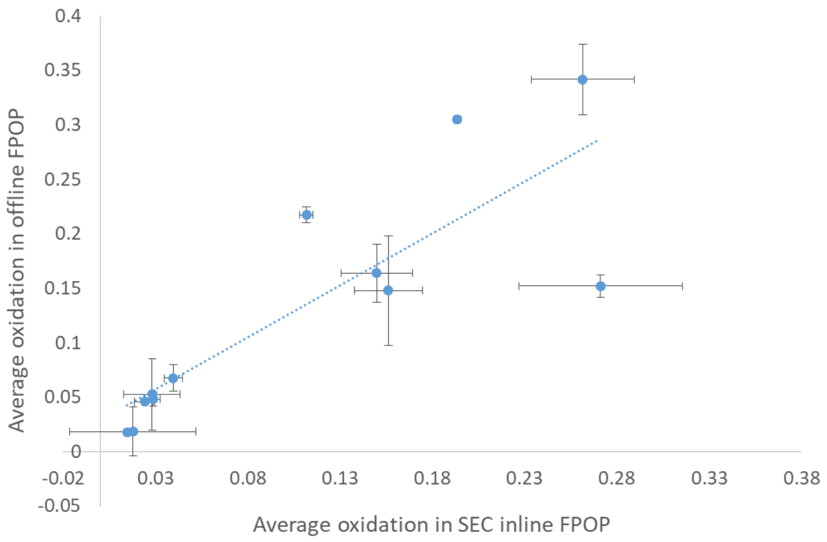


**Extended Data Fig. 4. Comparison of LC-FPOP footprint of monomeric adalimumab with previously published offline FPOP data.** LC-FPOP data of the protein eluting from 26-30 minutes gave a strong correlation (Spearman's ρ = 0.75) with traditional offline FPOP data of a different monomeric adalimumab sample previously published in July 2019[24]. The strong correlation in results indicate the ability of LC-FPOP to accurately provide HRPF data of conformers from mixtures.
